# Quantitative landscapes reveal trajectories of cell-state transitions associated with drug resistance in melanoma

**DOI:** 10.1101/2022.04.16.488373

**Authors:** Maalavika Pillai, Zihao Chen, Mohit Kumar Jolly, Chunhe Li

## Abstract

Drug resistance and tumor relapse in melanoma patients is attributed to a combination of genetic and non-genetic mechanisms. Non-genetic mechanisms of drug resistance commonly involve reversible changes in the cell-state or phenotype, i.e., alterations in molecular profiles that can help cells escape being killed by targeted therapeutics. In melanoma, one of the most common mechanisms of non-genetic resistance is dedifferentiation, which is characterized by loss of melanocytic markers. While various molecular attributes of de-differentiation have been identified, the transition dynamics remains poorly understood. Here, we construct cell-state transition landscapes, to quantify the stochastic dynamics driving phenotypic switch in melanoma based on its underlying regulatory network. These landscapes reveal the existence of multiple alternative paths to resistance – de-differentiation and transition to a hyper-pigmented phenotype. Finally, by visualizing the changes in the landscape during *in silico* molecular perturbations, we identify combinatorial strategies that can lead to the most optimal outcome – a landscape with the minimal occupancy of the two drug-resistant states. Therefore, we present these landscapes as platforms to screen possible therapeutic interventions in terms of their ability to lead to most favourable patient outcomes.

## Introduction

Phenotypic heterogeneity in melanoma has been attributed to drug resistance, relapse and recalcitrance in patients ^1–4^. Clinically, phenotypic classification of tumor samples was introduced to distinguish between the metastatic potential of melanoma tumors. Cells were classified as proliferative or invasive, based on their ability to contribute to tumor growth (non-metastatic) and their ability to migrate to secondary tumor sites (metastatic), respectively ^5^. These two phenotypes were shown to be capable of switching between one another and giving rise to heterogenous tumors independently ^6,7^. Although the classification started as a binary system for metastatic and nonmetastatic samples, intermediate phenotypes displaying features of both the extreme phenotypes have also been identified ^5,8–10^. A recent study classified melanoma samples into four phenotypes: melanocytic, transitory, neural crest stem cell like (NCSC) and undifferentiated (in a decreasing order of proliferative behavior, or in an increasing order of invasive traits) ^11^. Treatment of melanocytic samples with BRAF inhibitors (BRAFi) or MEK inhibitors (MEKi) can cause cells to follow a distinct dedifferentiation trajectory, where they first transition to being intermediate phenotypes on the proliferative-invasive axis (transitory and NCSC) and finally get undifferentiated ^12–14^. De-differentiation has been dynamically characterized as a response to targeted therapy, a response driven by underlying gene regulatory network that determines the changes in gene expression (during drug treatment or other perturbations) and the consequent switching of phenotypes. Dynamical simulations of such networks suggest that the four abovementioned phenotypes exist as “microstates” within two larger “macro-states”, namely the proliferative and invasive phenotypes ^15^. However, the stochastic cell-state transition dynamics among these four phenotypes remain to be determined.

Besides de-differentiation, recent studies have identified alternate pathways to therapy resistance ^16–18^. Single cell profiling of patient-derived xenograft models revealed an alternate resistance pathway where cells increase expression of pigmentation genes and are referred to as hyper-pigmented ^19^. The hyper-pigmented phenotype is characterized by high expression of melanocyte differentiation marker MITF and its downstream targets such as PMEL, TYR and MLANA ^20^. While the existence of such an alternate pathway has been characterized, the underlying regulatory mechanism and the cell-state transition trajectory to this phenotype remains to be deciphered yet.

The (co-) existence of multiple cell-states is visualized as switching among diverse “attractors” in a given potential landscape, thus enabling the emergence of non-genetic heterogeneity. Waddington’s landscape is a classic example of such multi-stable potential landscapes being used to explain the dynamics of differentiation of cells, where each differentiated cell type is a stable state. In the context of cancer biology, such landscapes have been used to explain progression from a normal to cancerous cell state, switching to a cancer stem-cell like state, and epithelial-to-mesenchymal transition in carcinomas ^21–26^.

Here, we explain the existence of categorical multi-stability (**Fig. 1A**) and dynamics governing resistant cell-fate decisions in melanoma, using quantitative landscapes. We infer cell-state transition probabilities and identify the de-differentiation trajectory followed by cells as the most likely transition path in the given landscape. Further, we reveal conditions under which cells can take an alternate trajectory for drug-resistance: the acquisition of hyper-pigmented state. Finally, we use these landscapes to identify combinatorial strategies to promote favorable outcomes. Our model not only provides a mechanistic understanding of phenotypic heterogeneity and underlying dynamics during phenotypic switching, but also acts as a platform for primary screening of target genes to identify potential therapeutic strategies that can drive desirable outcomes.

**Figure 1:**
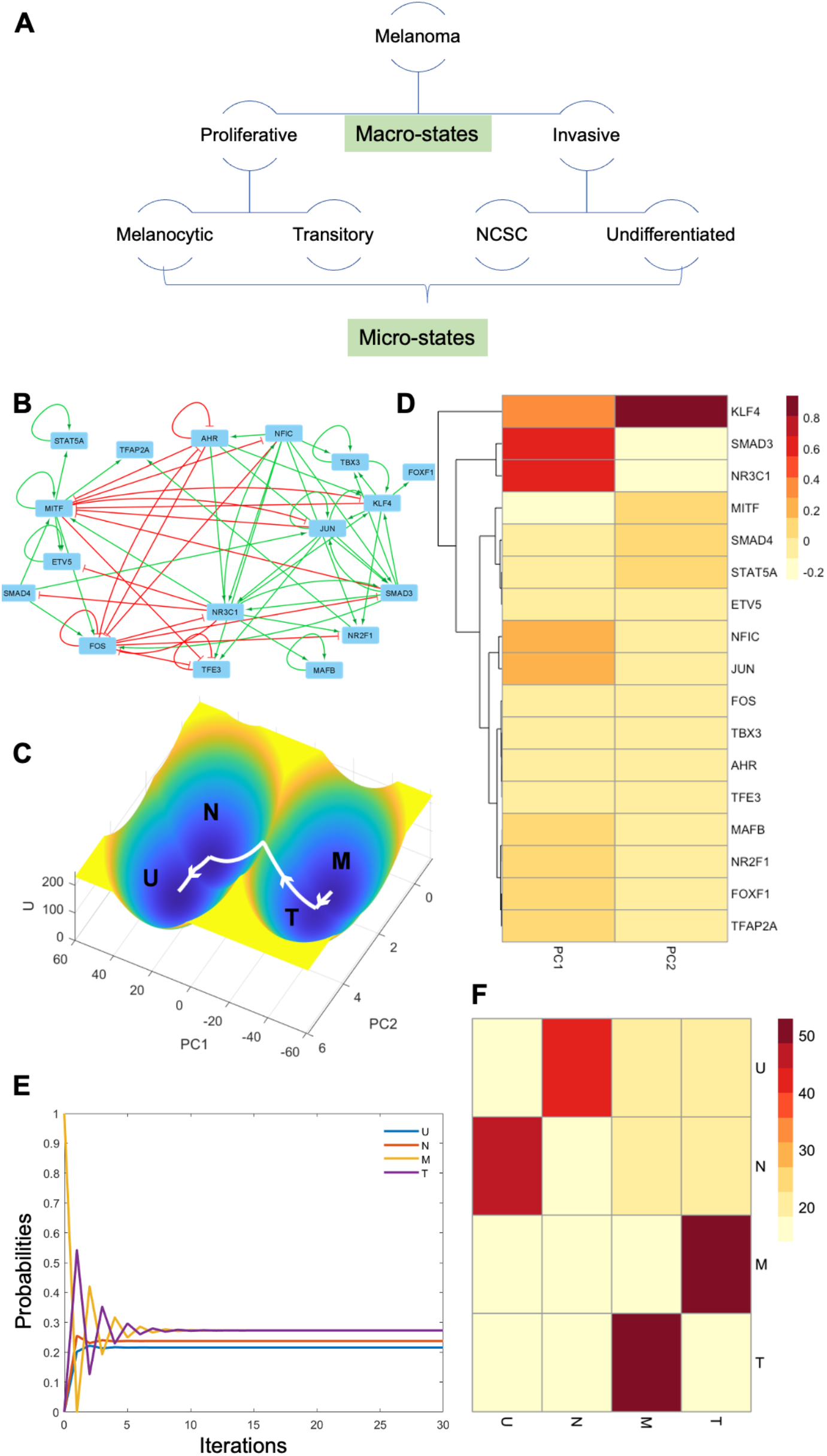
Energy landscapes explain the existence of categorical multi-stability in melanoma tumors. **A.** Schematic representation of categorical multi-stability. **B.** Gene regulatory network (GRN) governing phenotypic heterogeneity in melanoma. **C.** Energy landscape for parameter set giving rise to 4 stable states/ phenotypes. **D.** Heatmap for PCA loading coefficients categorize genes as proliferative and invasive. **E.** Changes in transition probabilities from Melanocytic phenotype with simulation time. **F.** Heatmap for transition probability matrix derived from energy landscape (X-axis: To; Y-axis: From). Color bar denotes percentage probabilities.

## Materials and methods

### RNAseq datasets used

scRNAseq dataset GSE115978 was used for pseudo-time analysis and measuring intra-tumoral heterogeneity ^27^. The samples were filtered to remove non-tumor cells and only cells labelled as malignant melanoma cells were considered for the analysis. Since the dataset comprised of several single cell samples from multiple tumors, it was suitable for quantifying intra-tumor as well as inter-tumor heterogeneity. ^10,19^

### Pseudo-time analysis

Pseudo-time analysis was done as previously reported ^15^. AUCell scores ^28^ for the proliferative, hyper-pigmented, NCSC and invasive phenotype gene sets ^19^ were used to identify trajectory of cells along the pseudo-time axis.

### Measuring heterogeneity

Intra-tumoral heterogeneity was measured using Shannon’s diversity Index (S) which is used to quantify the uncertainty or entropy of a system ^29^. It is calculated as:

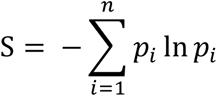

where, *p_i_* is the proportion of the *i^th^* phenotype in the population and *n* is the total number of phenotypes present.

### Landscape Generation

We use *X*(1), *X*(2),…, *X*(*n*) to describe the state variables of cells in the melanoma system, which represent gene expression levels. To obtain the potential landscape, we need to calculate the probability distribution of the system. The probability distribution of the system evolves with time and is recorded as *P*(*X*(1), *X*(2),…,*X*(*n*), *t*), which is governed by a diffusion equation. We follow a self-consistent mean field approach to split the probability into the products of the individual ones ^30–34^, i.e., 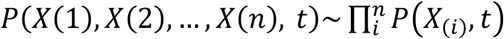 and solve the probability self-consistently.

Specifically, we use Gaussian approximation to calculate the probability distribution, which means that we need to calculate two moments, the mean and the variance. When the diffusion coefficient *D* (characterizing the noise level) is small, the moment equations can be approximated to ^34–36^:

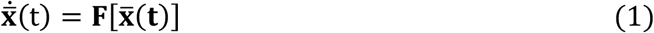

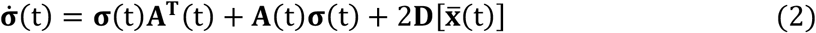

Here, **x, σ**(*t*), and **A**(*t*) are vectors and tensors, and **A^T^**(*t*) is the transpose of **A**(*t*). The elements of matrix *A* are specified as: 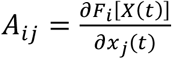.

We can solve 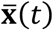 and **σ**(*t*) numerically. Here, we only consider the diagonal elements of **σ**(*t*) based on the mean field approximation. The probability distribution for each variable can be obtained from the Gaussian approximation as:

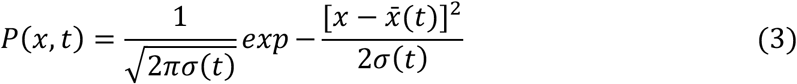

Here, 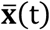 and **σ**(*t*) are the solutions of Eq. (1) and (2), and *P*(*x,t*) corresponds to the probability distribution of one attractor. The total probability of the system is the weighted sum of all these probability distributions. With the total probability, we can construct the potential landscape by *U*(*x*)=-*lnP_ss_*(*x*) ^30,34^, with *P_ss_* representing steady state probability distribution.

### Transition paths

A dynamical system in the fluctuating environments can be addressed by:

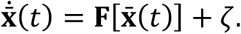

Here, **x** = (*x*_1_(*t*), *x*_2_(*t*),…, *x_n_*(*t*)) represents the vector of the gene expression level. **F**[**x**(*t*)] is the vector for the driving force for system, and *ζ* is the Gaussian white noise, which satisfying *E*[*ζ_i_*(*t*)*ζ_j_*(0)] = 2*Dδ_ij_ δ*(*t*), where *D* is the constant diffusion coefficient and

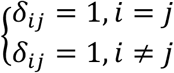

Following the approaches based on the Freidlin-Wentzell theory ^37^, the most probable transition path from attractor *i* at time 0 to attractor *j* at time T, can be acquired by minimizing the action functional over all possible paths:

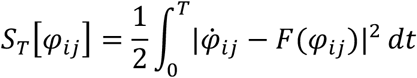

This path is called the minimum action path (MAP). We calculated MAPs numerically by applying minimum action methods used in previous study ^38^.

### Calculation of transition matrix

The transition action characterizes the probability of transition from one attractor state to another, i.e., smaller transition action corresponds to larger transition probability. Based on the results of transition actions between attractors, we can estimate corresponding transition probability according to *P_ij_* ∝ *e*^-*S_ij_*^, where *S_ij_* represents the transition action from attractor *i* to attractor *j*, while *P_ij_* represents the transition probability from attractor *i* to attractor *j*. In this way, we can obtain an estimated transition probability matrix among five stable states. Considering the state transition process among attractors as a Markov process, we can calculate the probability evolution of each attractor state given any initial probability distribution of five stable states.

## Results

### Quantitative landscapes capture categorical multi-stability in melanoma

We simulated the dynamics of a previously defined gene regulatory network (**Fig. 1B**) that explains the existence of four phenotypes, namely, Melanocytic (M), Transitory (T), NCSC (N) and Undifferentiated (U) ^15^, and generated corresponding landscape showcasing these four phenotypes observed in melanoma (**Fig. 1C**). Previously, in order to estimate parameters and simulate the network-derived model, we had used RACIPE (**RA**ndom **CI**rcuit **PE**rturbation), which generates a large number of models by using an ensemble of parameter sets such that kinetic parameters are sampled randomly from a distribution corresponding to biologically feasible *in vitro* values ^39,40^. For this study, we selected parameter sets corresponding to tetra-stable or penta-stable models (i.e., models that give rise to 4 or 5 stable states, respectively) (**Supplementary Table 3**).

In our earlier work ^15^, we had proposed that the underlying regulatory network gives rise to multi-stability comprising of two broader categories or “macro-states” (Proliferative and Invasive) that can be subdivided into 4 phenotypes or “micro-states” (Proliferative comprises of Melanocytic, Transitory; while Invasive comprises of NCSC and Undifferentiated) (**Fig. 1A**). However, this categorization was entirely based on clustering algorithms and Euclidean distance metrics applied to steady-state values obtained from RACIPE simulations. These methods do not provide any insight into the actual dynamics of the system. Thus, here, we focus on the relative stability of each state and its ability to transition to other states to quantify multi-stability transition dynamics in the system (characterized by the depth of the corresponding “attractor” on the landscape).

For the tetra-stable parameter sets, the landscape highlights the existence of these four phenotypes as four attractors lying within two larger “macro-states” (proliferative and invasive) (**Fig. 1C**). The melanocytic and transitory phenotypes lie within the proliferative macro-state and the NCSC and undifferentiated phenotypes lie in the invasive macro-state, as expected. Here the landscape is acquired based on a dimension reduction of landscape (DRL) approach ^21^, by projecting the highdimensional landscape to a two-dimensional space (PC1 and PC2). DRL identifies the contribution of each gene to the variability in the expression among the phenotypes. The loading coefficients corresponding to first two principal components displayed clear segregation of proliferative genes (MITF, FOS, STAT5A, ETV5, SMAD4) and invasive genes (KLF4, NR3C1, SMAD3, JUN, NFIC, AHR, TBX3) (**Fig. 1D, Table 1A**). We also observe that the most likely transition path of cells recapitulates the dedifferentiation trajectory (i.e., transition from M to T to N to U; highlighted using white arrows in Fig 1C). Transition probabilities for cells in the M state, display maximum likelihood for transition to the T phenotype (**Fig. 1E-F, Table 1B**). These results are captured in three separate parameter sets/models that were selected (**Fig. S1A-B, Supplementary Table 1**), thus suggesting more generic features of the underlying tetra-stable landscapes corresponding to melanoma.

### Penta-stability explains the existence of multiple trajectories for drug resistance

Besides de-differentiation, alternative trajectories to drug resistance in melanoma have been reported ^16,19^. The hyper-pigmented phenotype is a resistant phenotype that does not arise out of de-differentiation *per se*. It is characterized by an increase in melanocytic markers such as MITF, PMEL, TYR and MLANA which are commonly associated with pigmentation in melanocytes. We used our existing network to test whether it could explain the existence of this phenotype. To identify the fifth phenotype, we used k-means clustering (k=5) on the RACIPE simulation dataset that was previously generated ^15^. Parameter sets that gave steady state solutions for all five clusters were used to generate landscapes (**Supplementary Table 3B**).

The landscapes revealed that the fifth state, which is characterized by very high levels of MITF, fell outside the de-differentiation trajectory (highlighted using white arrows), suggesting that it represented the Hyperpigmented (H) phenotype (highlighted using pink arrows) as previously reported ^19^ (**Fig. 2A, Table 2A)**. The melanocytic and transitory phenotypes showed the highest probability of switching to the hyperpigmented phenotype (**Fig. 2B, Table 2B**). Similar results were witnessed for an alternate penta-stable parameter set too (**Fig. S2A-B, Supplementary Table 2**).

**Figure 2:**
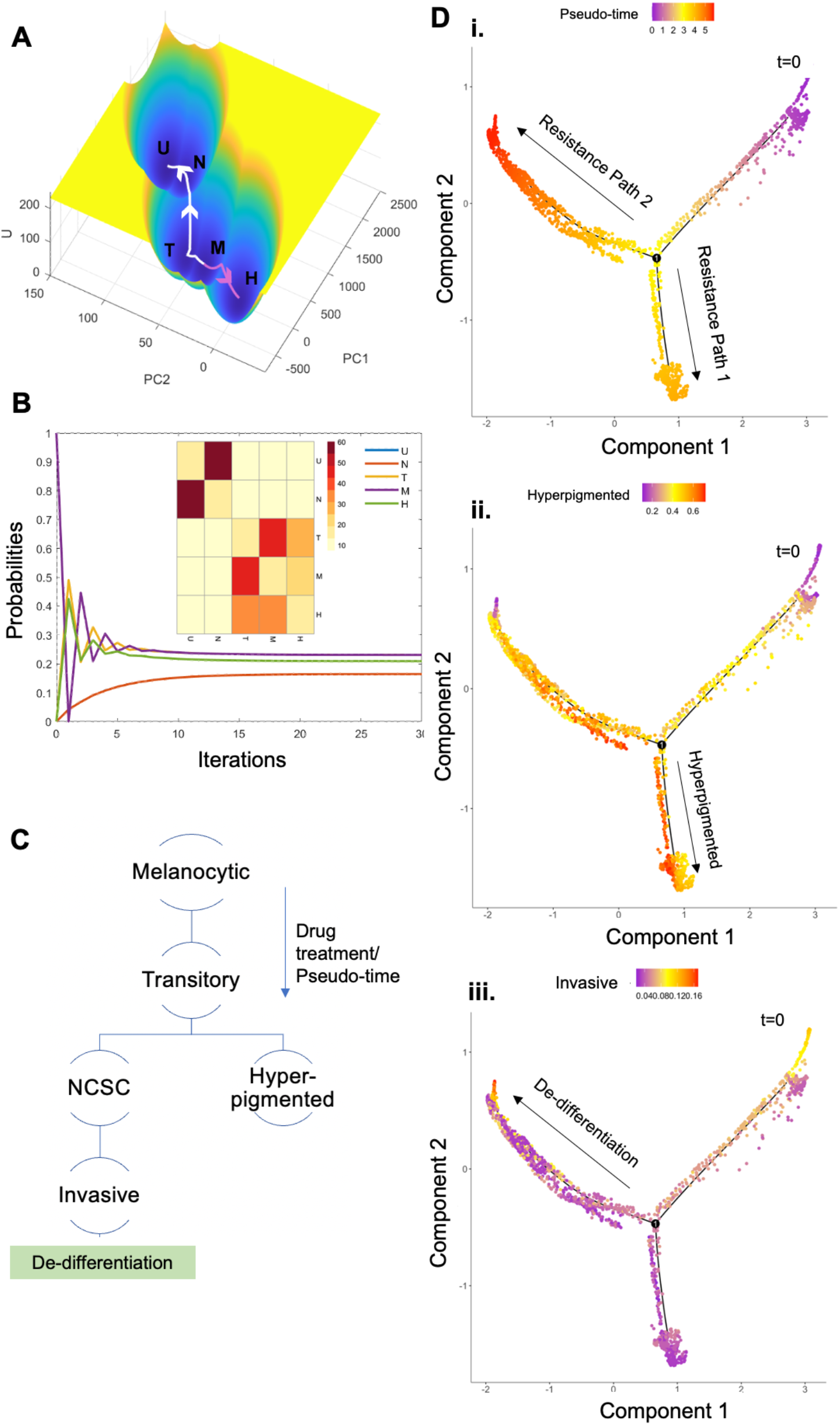
Energy landscapes explain the existence of multiple paths to drug resistance. **A.** Energy landscape for parameter set giving rise to 5 stable states/ phenotypes. **B.** Changes in transition probabilities from Melanocytic phenotype with simulation time. Inset represents heatmap for transition probability matrix derived from energy landscape (X-axis: To; Y-axis: From). Color bar denotes percentage probabilities. **C.** Schematic representation of multiple drug resistant trajectories. **D.** Pseudo-time analysis reveals two trajectories to drug resistance in GSE115978. **i**. Pseudo-time analysis. **ii**. AUCell scores for hyper-pigmented gene set. **iii.** AUCell scores for invasive gene set.

To validate this prediction made by landscape quantification, we performed pseudotime analysis of single-cell tumor samples (GSE115978). This analysis recapitulated similar trends, where melanoma cells transition, along the pseudo-time axis, from being in a proliferative phenotype to a transitory phenotype which further bifurcates to give rise to the de-differentiation and hyper-pigmentation trajectories for drug resistance (**Fig. 2C-D, S2C-D)**, thereby underlining two alternative or non-overlapping trajectories to resistance in melanoma. Scores for each phenotype were calculated using AUCell for gene sets reported earlier ^19,28^. The genes regulating the hyperpigmented phenotype (red labels) and the invasive phenotype (blue labels) form two mutually inhibiting ‘teams’ of genes (**Fig. S2E**), i.e., genes regulating a phenotype are positively correlated with each other and negatively correlated with genes regulating the other phenotype. Such ‘teams’-like interactions have been shown to play a role in regulating phenotype switching and multi-stability in biological systems ^41,42^. Transition paths based on the energy landscapes for the two trajectories (de-differentiation and hyperpigmentation) reveal the changes in gene expression as cells transition from one state to another (**Fig. S3)**. The state transitions during de-differentiation (M to T to N to U) reveal a clear switch from high levels of expression of proliferative and melanocytic markers (MITF, FOS, SMAD4, ETV5) to lower levels and a concomitant increase in invasive markers (SMAD3, AHR, NFIC, KLF4, JUN) (**Fig. S3A-B**). The state transitions for hyperpigmentation (M to H), not only reveal an increase in expression levels of MITF, a well characterized regulator of pigmentation in melanocytes, but also an increase in expression of ETV5 and TFAP2A. Previous studies have reported regulation of MITF, ETV5 and TFAP2A by super-enhancer motifs during hyper-pigmentation ^43^. While MITF and TFAP2A have been reported to display clear upregulation during hyperpigmentation. Overall, our model is able to recapitulate multiple paths to drug resistance and the associated transitions in cell states observed in melanoma tumors.

### Quantifying intra-tumor and inter-tumor heterogeneity in melanoma samples

After analyzing abovementioned experimental datasets, we investigated single-cell transcriptomic data gathered from multiple tumors. Interestingly, upon segregating these samples based on the tumor, we observed that most tumors comprised of a heterogenous population of cells, i.e., cells from a single tumor were proliferative, de-differentiated/invasive and/or hyper-pigmented (**Fig. S2F**), thus highlighting intratumor heterogeneity. The frequencies of these phenotypes also varied across tumors, thereby showcasing inter-tumor heterogeneity as well.

To quantify intra-tumor heterogeneity, we used the Shannon Diversity Index (SDI) (**Fig. 3A**). Higher SDI indicates higher levels of diversity and thereby higher uncertainty in determining the phenotype of a random cell. For tumors comprising of cells from a single phenotype, SDI is 0 (Mel121, Mel81, Mel84), and tumors with a high proportion of a single phenotype have SDI close to 0 (Mel106, Mel79, Mel80). Intriguingly, very few tumors had hyper-pigmented phenotype as the dominant phenotype, suggesting that this phenotype can be perhaps less abundant in tumors. This trend is also captured by the relative stability of the hyper-pigmented phenotype in simulated data (**Fig. 3B**). Only a small fraction (<25%) of solutions corresponded to the hyperpigmented phenotype in both cases. This low abundance might have possibly contributed to this phenotype being discovered more recently under the lens of single cell profiling ^19^, as opposed to previous bulk-level studies on heterogeneity in melanoma ^11^.

**Figure 3:**
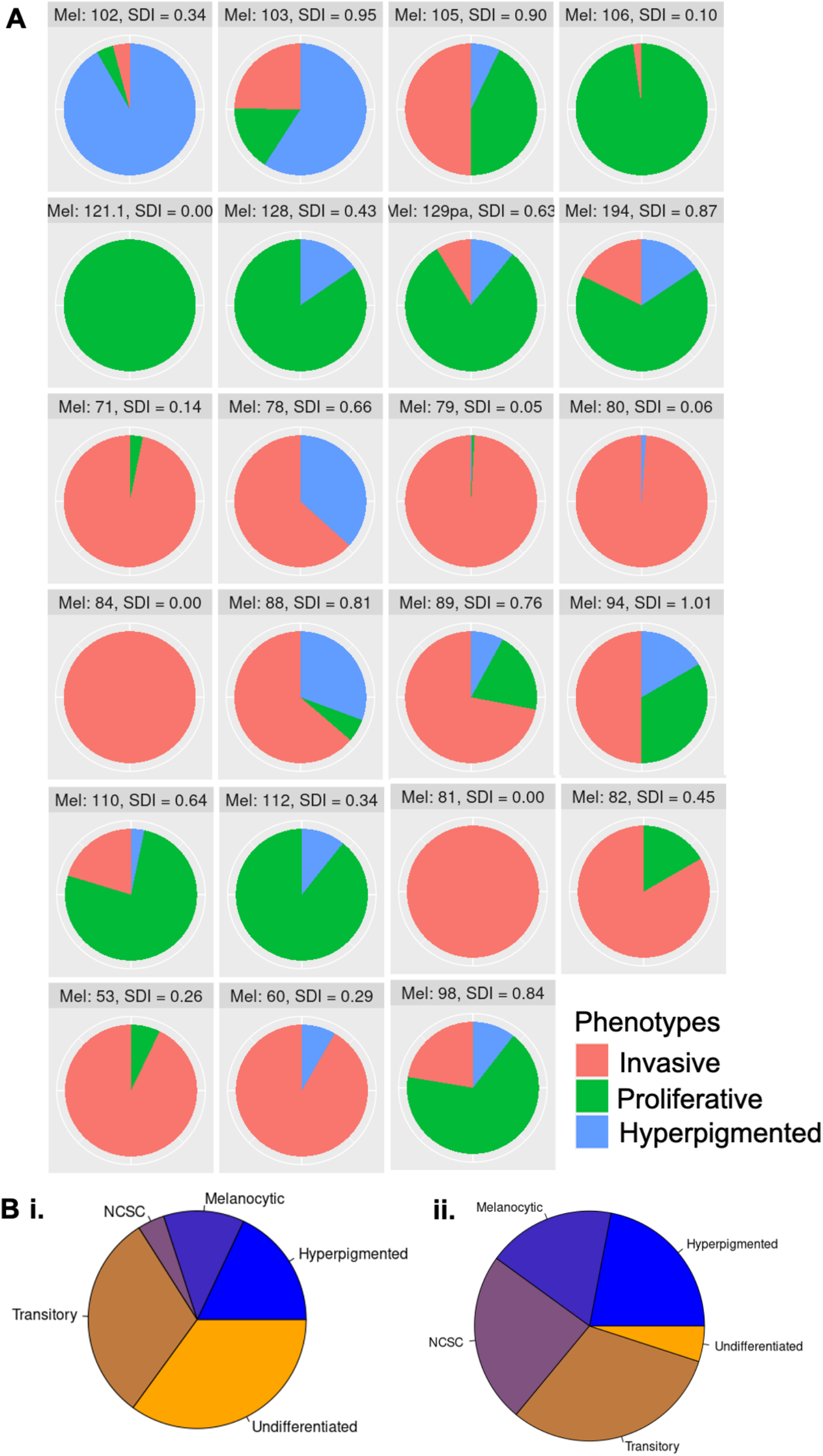
Patterns of intra-tumoral heterogeneity vary across patients. **A.** Pie charts representing proportion of cells belonging to each phenotype in a tumor. Titles include tumor name and Shannon’s Diversity Index (SDI) to quantify ITH. **B.** Pie charts representing proportion of initial conditions giving rise to each of the 5 steady states/ phenotypes in simulations for 2 penta-stable parameter sets.

### *In silico* knockdowns predict optimal drug treatment strategy

To check whether our landscape-based model could recapitulate the effects of targeted therapy as noted experimentally, we knocked down (KD) *in silico* some commonly targeted genes. We used MITF KD to mimic the effect of BRAF inhibition, since BRAF is a gene upstream of MITF ^44^. A recent study also highlighted the ability to overcome drug resistant phenotypes by knocking down SMAD3 or its upstream regulator AHR, both of which are included in our network ^45^. Knockdown of MITF alone, led to a loss of the hyper-pigmented phenotype from the landscape (as compared to the unperturbed landscape), leaving only the proliferative (melanocytic and transitory) and de-differentiated phenotypes (NCSC and Undifferentiated) (**Fig. 4A-B**). SMAD3 KD led to the loss of invasive and de-differentiated phenotypes, leaving only the hyperpigmented and proliferative ones (**Fig. 4C**). A recent study identified AHR and SMAD3 as regulators of resistant phenotypes that arise during BRAFi induced dedifferentiation. Knocking down AHR or SMAD3 prevented the emergence of resistant phenotypes during BRAFi treatment ^45^. We recapitulated this experiment *in silico* by knocking down MITF and SMAD3 simultaneously. This led to the loss of both the dedifferentiated (U) and the hyper-pigmented (H) resistant states (**Fig. 4D**). Similar trends were also seen for combined knock down of AHR and MITF, further validating the ability of our model to make predictive outcomes for targeted therapy (**Fig. S4A**).

**Figure 4:**
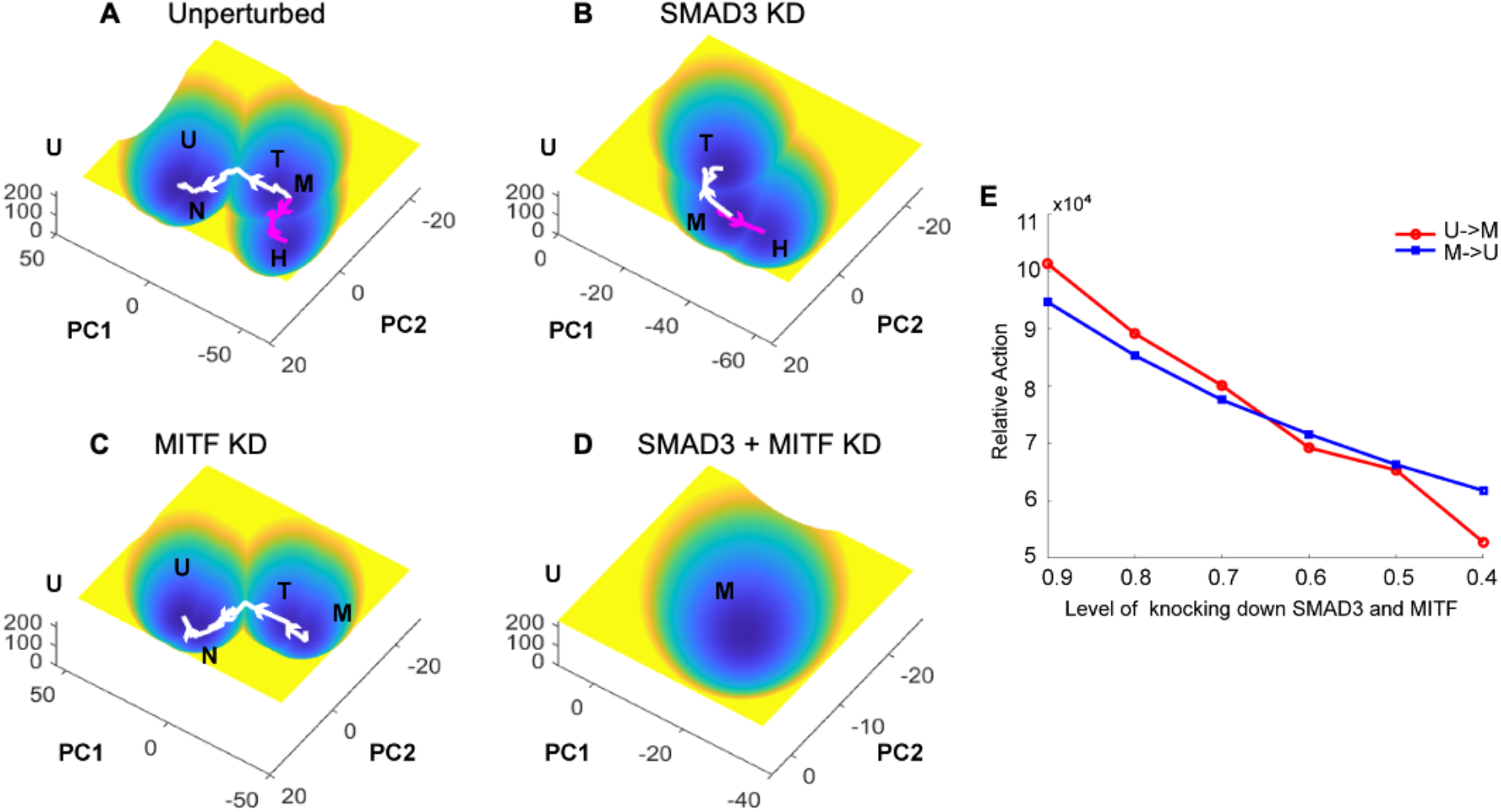
Energy landscapes recapitulate change in phenotypic distribution during targeted gene knockdown. **A.** Energy landscape for unperturbed system. **B.** Energy landscape for SMAD3 knockdown. **C.** Energy landscape for MITF knockdown. **D.** Energy landscape for MITF and SMAD3 dual knockdown. **E.** Variation in transition probability between melanocytic and undifferentiated phenotype with increasing knockdown effect on MITF and SMAD3.

Moreover, with an increase in the extent of knockdown of MITF and SMAD3, we observed a lower rate of de-differentiation and a higher rate of differentiation towards the melanocytic phenotype (i.e., the transition rate of dedifferentiation from the melanocytic to undifferentiated phenotype became lesser than the reverse transition) (**Fig. 4E**). Given the potential of our model, we tried various combinations of gene targets to identify optimal combinations of target genes that can give rise to a landscape lacking the resistant phenotypes (similar to SMAD3+MITF KD or AHR+MITF KD) (**Table 3, Table 4**). For instance, AHR and NFIC KD displayed a loss of invasive and hyperpigmented resistant phenotypes, but still maintained the transitory phenotype (**Fig. S4B**). AHR KD alone, also gave similar results (**Fig. S4C**) while NFIC KD alone led to the loss of the invasive phenotypes alone (**Fig. S4D**).

## Discussion

Phenotypic heterogeneity in melanoma has been identified as one of the major drivers of resistance to targeted therapy in melanoma. While previous studies have identified networks and mechanisms for how phenotypic heterogeneity is regulated ^10,15,46^, here, we present a dynamical perspective of underlying energy landscapes that determine the resistant fate and trajectories of a cell that undergoes molecular perturbations. These landscapes confirmed the existence of multi-stability as previously suggested ^15^ and capture the bifurcation of drug resistance trajectory into two distinct branches, as observed in time course experiments ^19^. By integrating data coming from *in silico* simulations and *in vitro* experiments, we also offer possible reasons for reduced abundance of hyper-differentiated phenotype in a tumor. Besides providing useful information on dynamics of cell-state transitions, these (pseudo-)potential landscapes also recapitulate the effects of various targeted therapies (KD of individual genes). Thus, they provide a basis for running high-throughput screening of target gene combinations to identify therapeutic strategies to overcome drug resistance in cells.

Multi-stability was proposed to underlie the existence of multiple levels of classification of phenotypic heterogeneity in melanoma. We previously suggested that the initial system of classification of samples as proliferative and invasive could further be resolved into four phenotypes: Melanocytic, Transitory, NCSC and Undifferentiated. Previously, this analysis was restricted to using distance-based metrics that provided no insight into transition rates and cellular dynamics. Energy landscapes are able to reveal this property of phenotypic classification by using potential (U) as a metric that quantifies the ability of cells in one state to transition to another ^34,47^. Although our model is only able to resolve up to five phenotypes, by increasing the resolution (by adding more genes or sub-networks for each phenotype to the existing 17-node network), possibly by adding sub-networks to the existing network, it might be possible to further resolve each of the 4 phenotypes into ‘micro-states’. Existence of such (often un-observable) ‘micro-states’ within larger ‘macro-states’ has been observed previously in the context of EMT as well ^48^. The presence of a continuum of heterogeneity can potentially be attributed to the existence of such micro-states ^46,47^.

Potential (U) measures the stability of a cell at each point on the landscape. It provides information on the probability of a cell in a given state to switch to another state. Higher potential barriers represent a lower likelihood of transitions between two states. These landscapes can be used to estimate the amount of “force” or perturbation needed to induce a switch from one attractor state to another. This can be used to identify molecular variables that can increase the transition probability towards a preferable state, therefore allowing us to push a population of cells towards a preferred state. This unleashes endless possibilities of tweaking the system to obtain desirable outcomes or predicting the effects of such perturbations on the outcome. Such (pseudo-) potential landscapes have been used extensively in developmental biology from modeling to understand cell-fate decision making and differentiation and can be useful tools to uncover cellular dynamics during cancer progression ^34,49,50^.

Our study delves into the effect of multiple drug-resistant phenotypes on intra-tumoral heterogeneity (ITH). We observe that the hyperpigmented phenotype forms a very small fraction of the total tumor population, at least in the 23 tumors investigated here in which only 2 tumors exhibited it as the dominant phenotype. Similar results (reduced abundance) were seen in our simulation results too, possibly explaining its rather recent identification/discovery, relative to that of four other melanoma phenotypes ^19^. At a bulk level, it is likely that the transcriptomic signature of this phenotype is masked. To quantify ITH, we used Shannon’s diversity index, an entropy-based measure of uncertainty in predicting the phenotype of a random cell ^29^. Interestingly, no consistent patterns were found among the multiple tumors, suggesting that variations at a genetic, microenvironmental or epigenetic level might influence the landscape and therefore the overall composition of the tumor ^51^. Differences in treatment strategies can also lead to different outcomes in tumor heterogeneity. The dataset used here (GSE115978) comprises of samples from treatment naïve patients and patients that have been treated with immune checkpoint inhibitors, which might also contribute to additional variations in ITH ^27^.

Here, we highlight the ability of landscape models to be used as a basis for predictive modelling of targeted therapy strategies. Conventional drug screening for all possible combinations is time- and resource-intensive. Thus, by using such predictive models, we can identify rather quickly optimal combinations of target genes to achieve durable outcomes, making it a potential tool for primary screening of possible treatment strategies. Moreover, the model can be expanded to suggest sequential targeting too.

## Acknowledgement

This work was supported by Ramanujan Fellowship awarded to MKJ by Science and Engineering Research Board (SERB), Department of Science and Technology (DST), Government of India (SB/S2/RJN-049/2018) and by Infosys Young Investigator award to MKJ supported by Infosys Foundation, Bangalore. MP is supported by KVPY fellowship (DST). C.L. is supported by the National Key R&D Program of China (2019YFA0709502) and the National Natural Science Foundation of China (12171102).

## Author contributions

MP and ZC performed research, analyzed data and wrote the first draft of the manuscript. MKJ and CL conceived and supervised research and worked on manuscript revisions.

## Data and Code Availability

All codes used in this manuscript can be found at https://github.com/csbBSSE/Melanoma_Landscape

## Supplementary Figures

**Figure S1:**
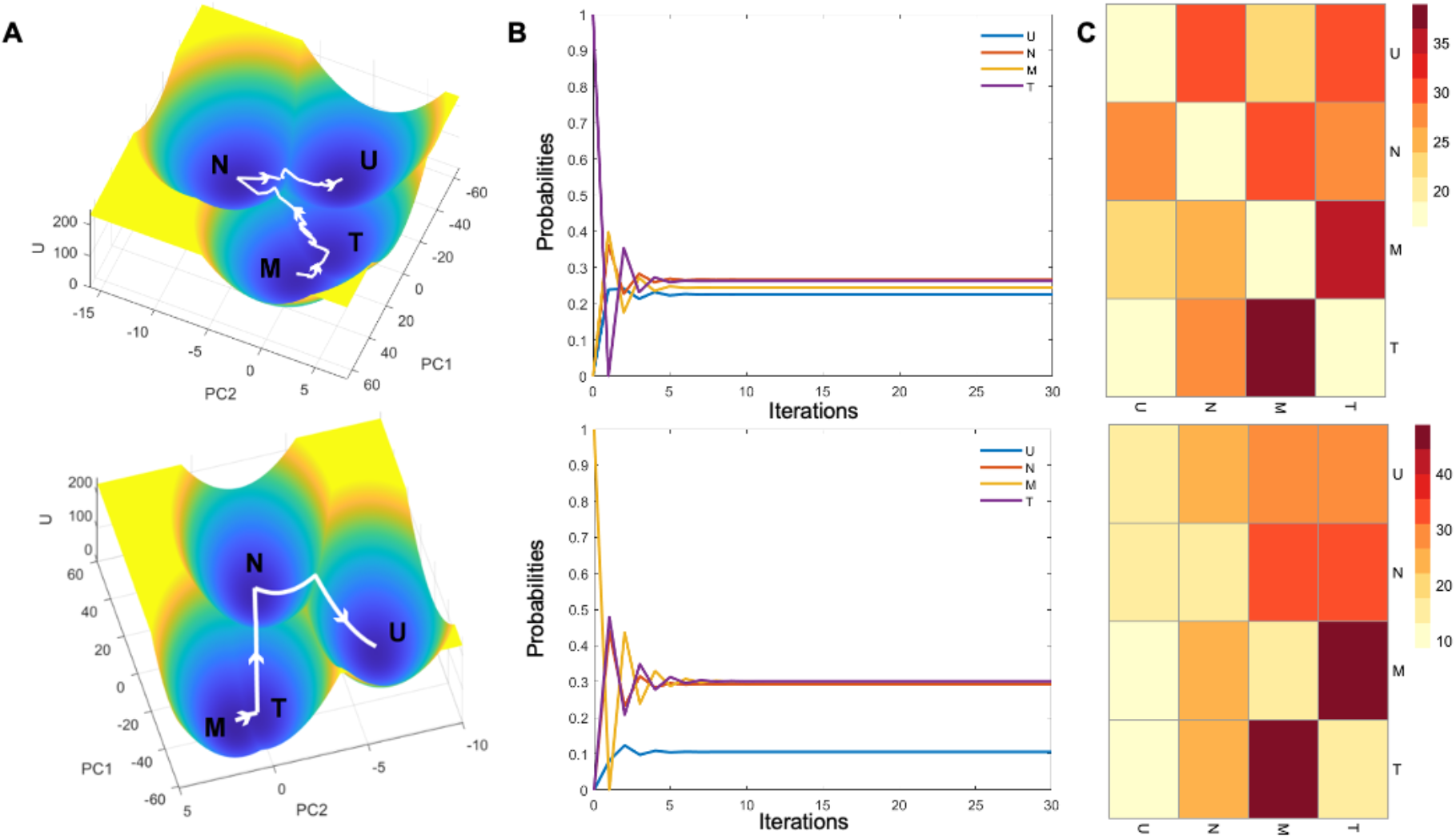
Energy landscapes for multiple tetra-stable parameter sets. **A.** Energy landscape for 2 parameter sets giving rise to 4 stable states/ phenotypes. **B.** Changes in transition probabilities from Melanocytic phenotype with simulation time. **C.** Heatmap for transition probability matrix derived from energy landscape. Color bar denotes percentage probabilities.

**Figure S2:**
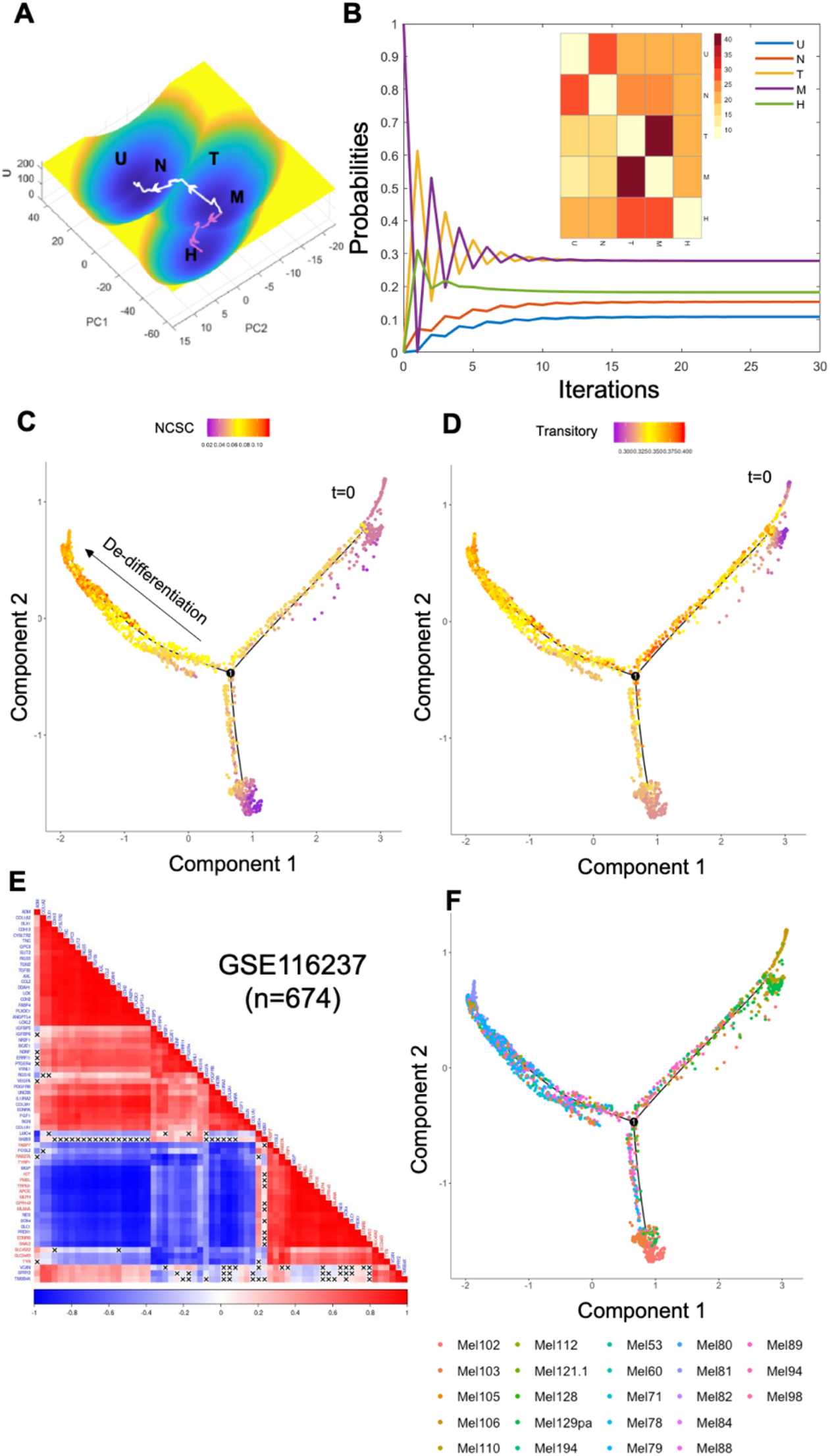
Melanoma tumor cells can take multiple trajectories to drug resistance. **A.** Energy landscape for parameter set giving rise to 5 stable states/ phenotypes. **B.** Changes in transition probabilities from Melanocytic phenotype with simulation time. Inset represents heatmap for transition probability matrix derived from energy landscape. Color bar denotes percentage probabilities. **C.** Pseudo-time analysis and AUCell scores for NCSC gene set. **D.** Pseudo-time analysis and AUCell scores for transitory gene set. **E.** Correlation plot for genes in hyper-pigmented geneset (Red labels) and Invasive gene set (Blue labels) for two ‘teams’ of players. Color bar denotes Spearman’s correlation coefficient. Crosses denote p>0.05. **F.** Segregation of single cell samples based on tumor of origin reveals intra-tumoral heterogeneity.

**Figure S3:**
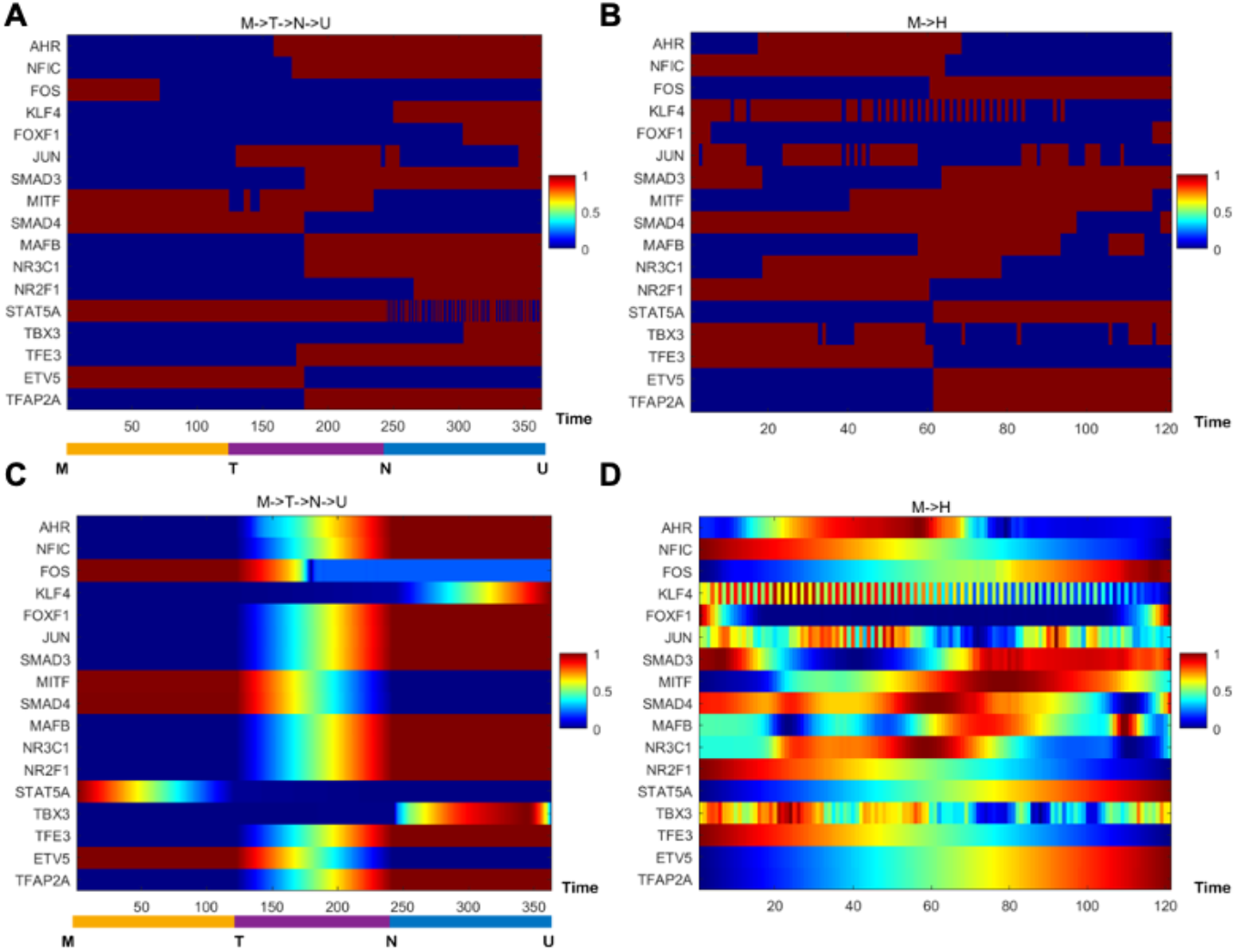
Transition paths in high-dimensional space for the two trajectories. Discretized expression during **A.** de-differentiation (M -> T->N -> U) and **B.** hyperpigmentation (M ->H). Continuous expression for transition path during **C.** de-differentiation (M -> T->N -> U) and **D.** hyper-pigmentation (M ->H).

**Figure S4:**
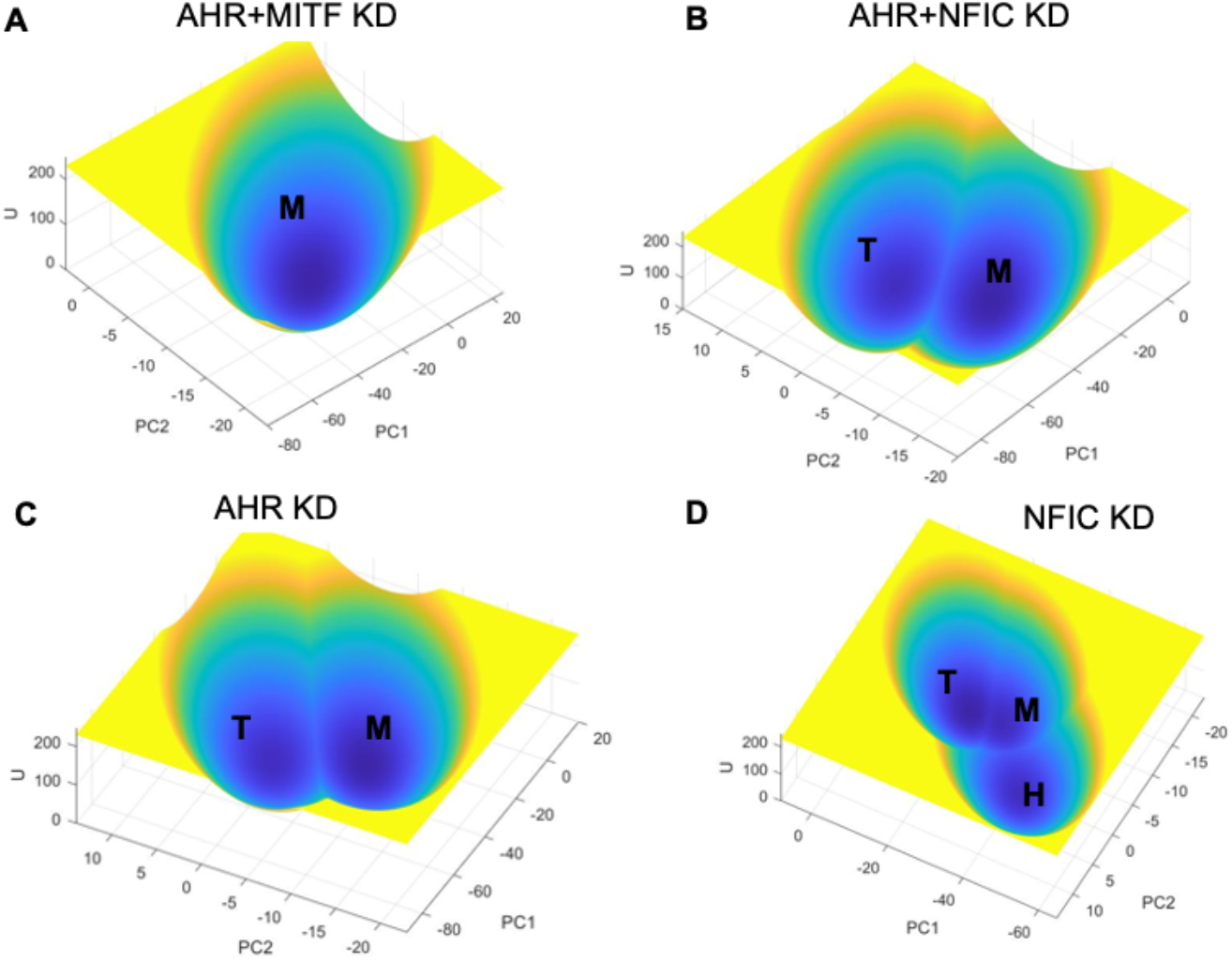
Energy landscapes provide a platform for screening target gene combinations. **A.** Energy landscape for AHR and MITF dual knockdown. **B.** Energy landscape for AHR and NFIC dual knockdown. **C.** Energy landscape for AHR knockdown. **D.** Energy landscape for NFIC knockdown.

